# Dietary restriction extends lifespan across different temperatures in the fly

**DOI:** 10.1101/2023.10.13.562222

**Authors:** Eleanor J Phillips, Mirre J P Simons

## Abstract

Dietary restriction (DR) has been consistently shown to extend lifespan across a range of taxa. However, it has recently been reported that DR does not extend lifespan at certain, namely lower, temperatures in flies (*Drosophila melanogaster*). Similar to the interpretation of other findings that appear to question DR’s universality, this finding has been interpreted as being an artefact of benign laboratory conditions. Here we re-test this hypothesis, now using a strain that shows robust lifespan extension at 25°C (across three prior experiments), and using a range of 5 diets across two temperatures, 18°C and 21°C. We found the DR longevity response to be robust, extending lifespan irrespective of temperature. We measured fecundity as a positive control for the DR phenotype, and found, as predicted, that DR reduces egg laying. It will be important for results that question DR as a phenotype to not be overinterpreted readily, as with a substantially larger sample size and a larger range of diets we were unable to replicate this prior work. Differences in experimental setup, genetic lines used and variation in the diet-lifespan reaction norm we suggest are responsible for this discrepancy. In addition, starting with a strain and conditions that show a lifespan extension by DR, as we do here, and then changing environment and/or genotype promises a more robust test of DR modulating factors.

## Introduction

Dietary restriction (DR) is one of the oldest known and best replicated life extending treatments in animals (Nakagawa et al., 2012; Simons et al., 2013). However, both its physiology and evolutionary biology remain hotly debated (Adler & Bonduriansky, 2014; McCracken, Adams, et al., 2020; Moatt et al., 2020; Piper et al., 2023). Given our incomplete understanding of DR and the intensity of study it continues to receive, it is perhaps unsurprising that with certain regularity studies report that DR is not extending lifespan and attribute this to certain circumstances, albeit genetic or environmental (Harper et al., 2006; Ja et al., 2009; Liao et al., 2010). Such studies can be interesting, but can also be distracting to the field if overinterpreted.

We recently argued that an absence of a DR response can be due to shift in reaction norm rather than a change in the dose-response (Simons & Dobson, 2023). Moreover, DR can arguably only be interpreted as absent when in the same study routine conditions result in a DR phenotype. A recent example of DR not extending lifespan of flies (*Drosophila melanogaster*) at lower temperatures has been interpreted as DR being a lab artefact with low temperature interpreted as representing stressful conditions (Zajitschek et al., 2023). There are several questions that can be posed to this study, for example: why did DR not extend lifespan at the most regularly used lab temperature of 25°C? Why did lifespan become truncated at low temperatures under DR? Why, if low temperatures induce stress, did flies show high levels of fecundity at those temperatures?

These questions largely involve interpretation of these findings. We also sought to question whether a strain that in our hands shows consistent and robust lifespan extension at DR at the arguably standard temperature of 25°C, showed no such response at lower temperatures (18°C and 21°C) using a range of five diets. We suggest the most reasonable test of the hypothesis posited by this recent work questioning DR, is to start with a strain that reliably and repeatedly shows DR under standard temperatures to then test whether lower temperatures negate the DR response. We use substantially larger sample sizes per treatment group (*N* is between 313 and 487 females) compared to the work that led to this hypothesis (*N* = 100) (Zajitschek et al., 2023).

We find that DR extends lifespan consistently across three separate experiments at 25 °C. We further find that DR extends lifespan irrespective of temperature (18 °C and 21 °C) and that its effect is highly quantitatively similar across temperatures even though lower temperatures, as expected, increased lifespan substantially.

## Results

First we wanted to confirm that we used a strain responsive to DR at the lab standard 25°C. In a strain (ywR) we have used extensively before (Drake & Simons, 2023; Gautrey & Simons, 2022) we found a significant DR longevity response in three other separate (before unpublished) experiments we previously conducted (P < 0.0001; Table 1). These experiments compared 2% yeast (our standard DR diet) to 8% yeast (the standard *ad libitum* diet used by our lab) (McCracken, Buckle, et al., 2020) (Figure 1). The response to DR varied slightly but significantly across these experiments (Chisq = 22.3, df=2, P < 0.0001, Table 1).

**Table 1.** DR extended lifespan across three different experiments in ywR flies. Log hazard estimates reported from separate models for each experiment as we found slight differences in the response to DR across these experiments, which is expected (Simons & Dobson, 2023).

| Experiment | logHR of DR (s.e.) | P | N |
| --- | --- | --- | --- |
| A | -2.68 (0.07) | < 0.0001 | 2,420 |
| B | -2.04 (0.13) | < 0.0001 | 566 |
| C | -2.36 (0.09) | < 0.0001 | 934 |

**Figure 1:**
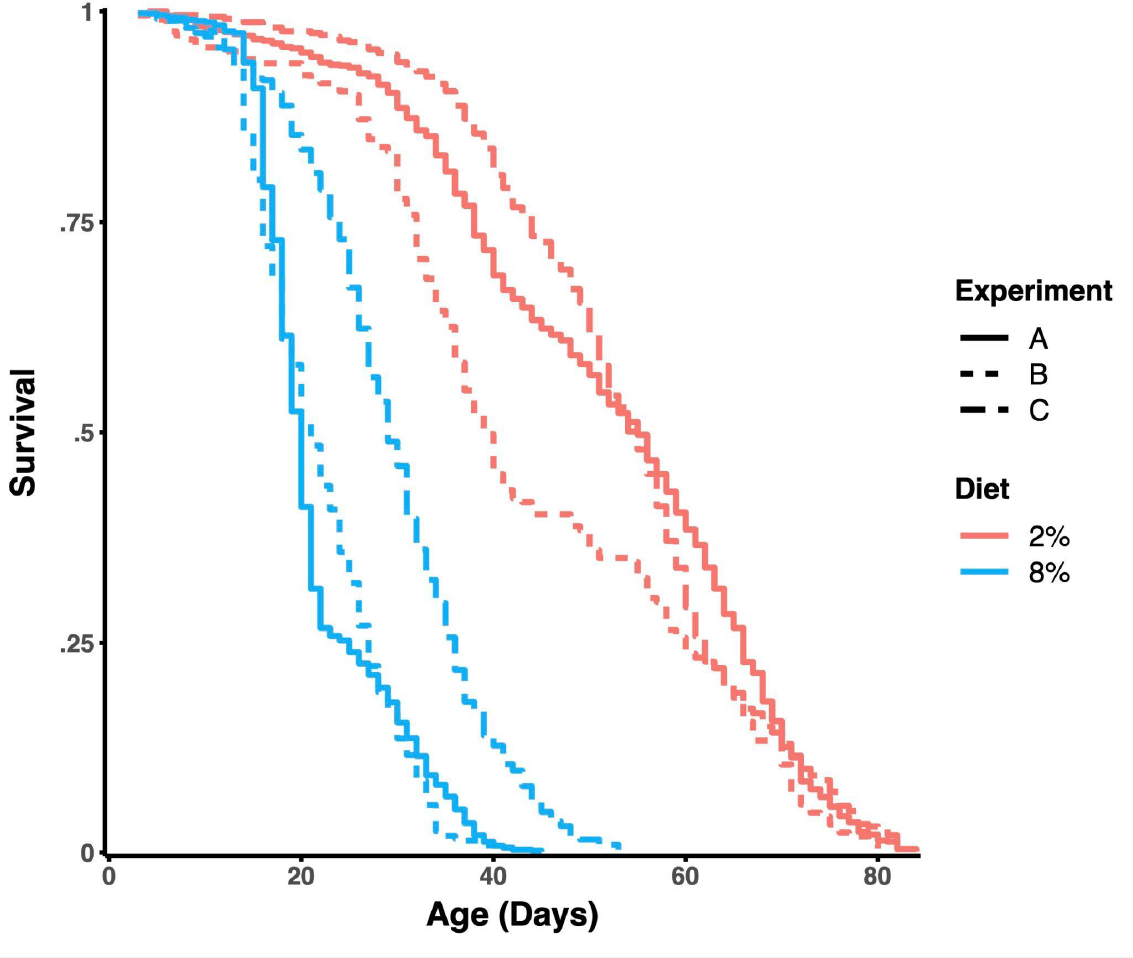
(a)Kaplan-Meier plot combining survival curves of three separate experiments, each using ywR flies on 8% and 2% yeast diets and kept at 25°C (N = 3,920 females total).

We then took this same strain and tested the DR response across five diets (2%, 4%, 6%, 8% and 10% yeast, keeping all other ingredients the same) at two temperatures 18 °C and 21 °C. We found the DR longevity response across both temperatures (P < 0.0001, Figure 2). Furthermore, there was no evidence that the diet effect was influenced by temperature (Chisq = 1.83, df=4, P = 0.77), and effects of diet were largely similar (Table 2, Figure 2a & 2b).

**Table 2.**
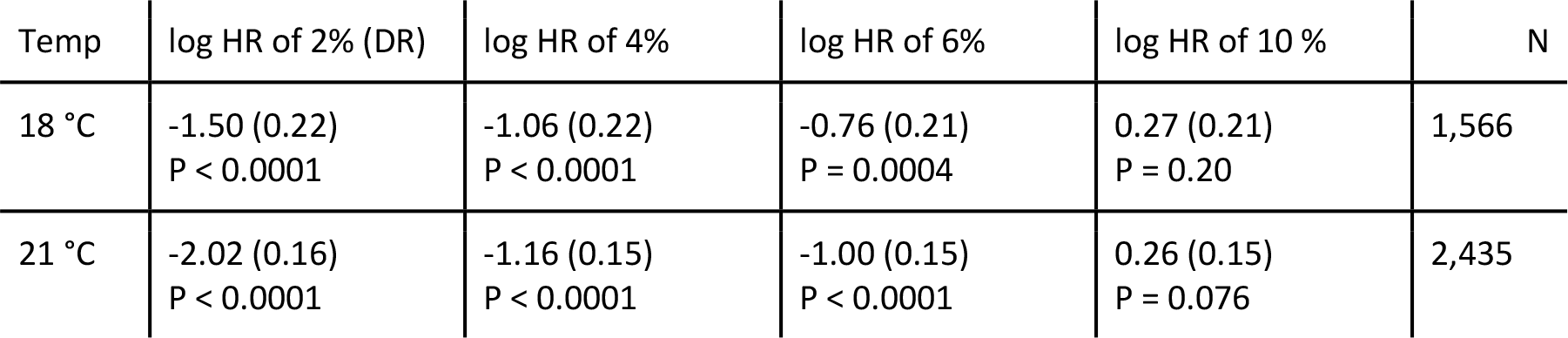
DR extends lifespan across both temperatures in ywR flies. Log hazard estimates are reported compared to the 8% yeast ad libitum diet (reference category). Estimates are largely the same and there is no evidence that the full diet response is modulated substantially across both temperatures. For 2% only, estimates are strongest at 25°C (Table 1), and decline slightly with decreasing temperature, but are still highly significant and strong. In linear hazard terms, risk is still lowered by 4.5 fold, and this resulted in an increase in median lifespan to 98 days at 2% yeast from 68 days at 8% yeast at 18°C (Figure 2).

**Figure 2:**
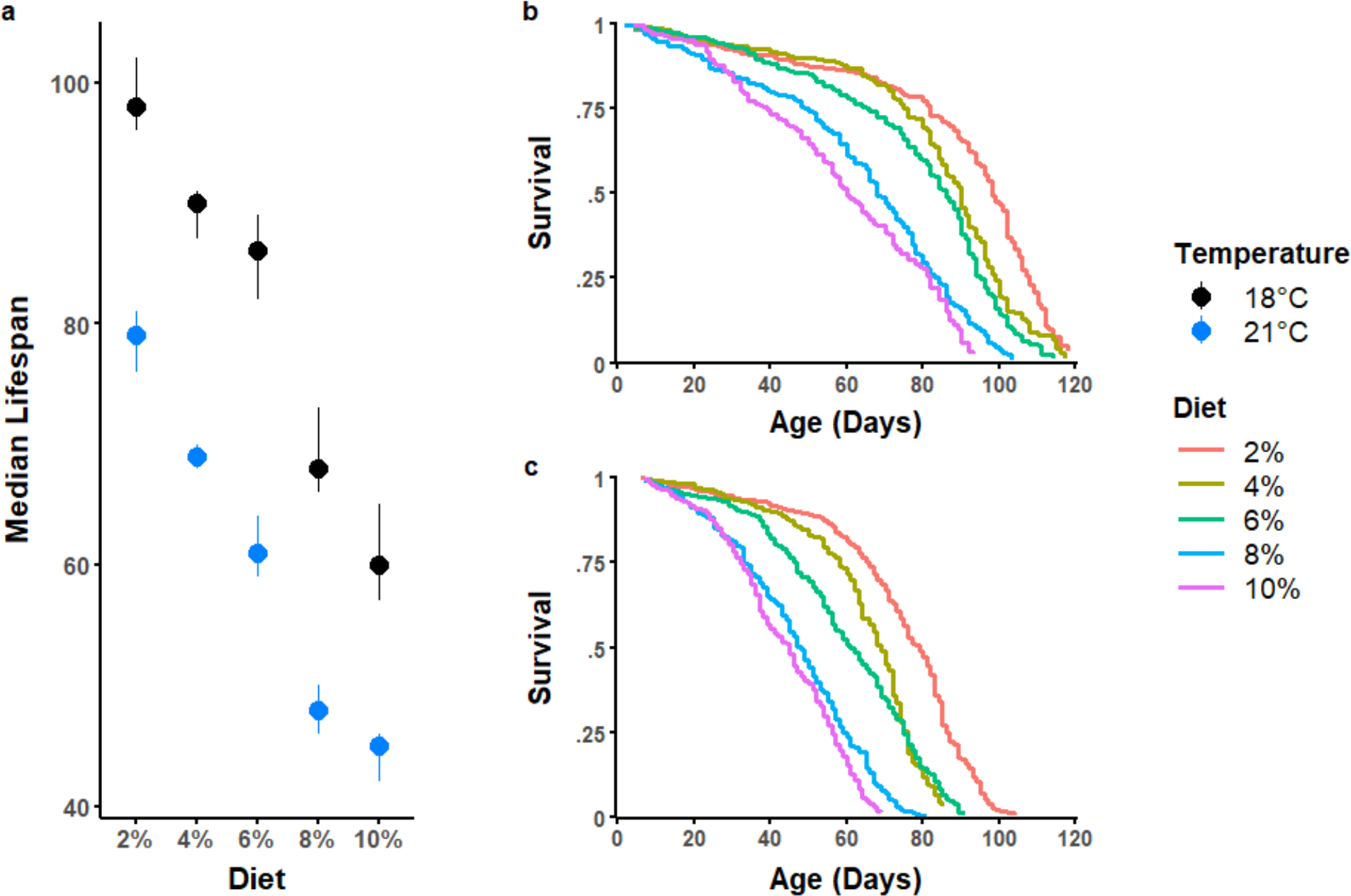
(a) Median lifespan and 95% CIs of yw drosophila fed diets with a range of yeast concentrations, kept at 18°C or 21°C. Respective Kaplan-Meier plots, showing survival rates of the different diet treatment groups at (b) 18°C or (c) 21°C. N = 4,063 females total; 297 to 501 per diet.

DR is classically associated with a decline in reproduction and is often used as a convenient readout to distinguish between a rescue from overfeeding and a true DR response (Gautrey & Simons, 2022; Grandison et al., 2009; McCracken, Buckle, et al., 2020). For this reason we measured egg laying at two time points during middle age (ages 36-41 to 40-45 days). We found no interaction between diet and temperature at either time point (P = 0.33 — 0.76) nor a main effect of temperature (P = 0.08 — 0.94). Higher yeast concentrations were associated with higher egg laying (P < 0.0001) and at the lowest yeast diet, on which flies also lived the longest (Figure 2), flies laid the fewest eggs (Figure 3).

**Figure 3:**
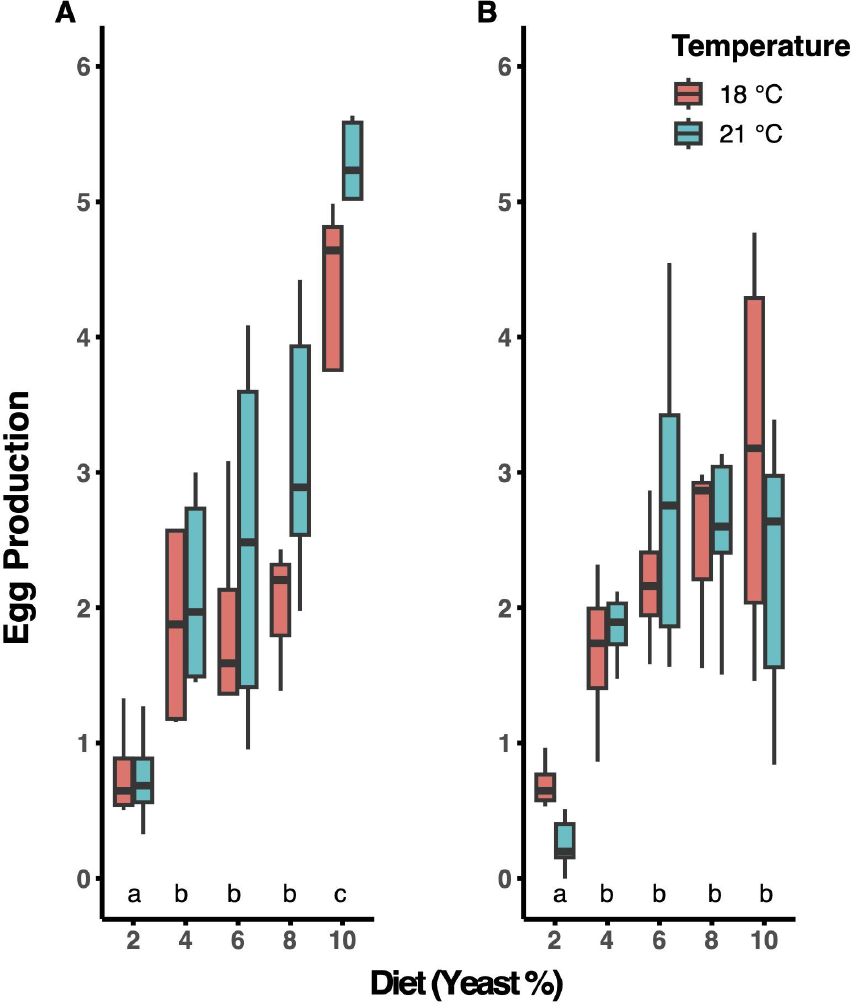
Mean egg production per fly across two days on different yeast diets, at two different time points (Mean ages of (A) 39 days and (B) 43 days). N = 49 demography cages respectively per time point. Letters (a,b,c) indicate significant differences based on post-hoc t-tests.

## Discussion

Our results unambiguously reinforce that DR can reliably increase lifespan in flies at standard lab conditions across different temperatures. As such our work fits with earlier work that used a combination of life-extending treatments, including temperature and DR, and found them to be largely additive (Kim et al., 2020; Shaposhnikov et al., 2022). Our findings and this prior literature contradict those of a recent study (Zajitschek et al., 2023) and counter their argument that DR constitutes a lab artefact.

These authors (Zajitschek et al., 2023) considered 25°C to be a low temperature for flies as it is lower than the ambient temperature in the climate of origin for the species. However, most experimental lines of flies will have been inbred or bottlenecked for many years under laboratory conditions, often at 25 °C. The precise original climate of the specific strain used is thus unlikely to still be physiologically relevant. Moreover, genetic variation inherent in this ‘outbred’ line can lead to unexpected variation in the population reaction norm to diet if genotypes present within the stock have different dose-responses to diet (Simons and Dobson, 2023). Furthermore, it is a potential confound that this prior study did not report a clear DR effect at the temperature flies are commonly grown at and the temperature at which DR is mostly studied, namely 25 °C.

Notably, heatshock protein expression is present at benign temperatures (25 °C) in *Drosophila*, with responses differing between lines (Bettencourt et al., 1999) and species (Kristensen et al., 2002; Sørensen et al., 2019). It is unclear therefore why lower temperatures should be interpreted as stressful. Especially as low temperatures increase lifespan (Mair et al., 2003) and at very low temperatures diapause is induced which results in mortality amnesia. Flies that are put at 25°C after a period in diapause resume their mortality trajectory as if they were in suspended animation (Tatar & Yin, 2001). In Zajitschek et al. 2023, it is further reported that egg laying is highest at relatively lower temperatures, again suggesting that these temperatures are not stressful, especially because these are also at the temperatures flies lived the longest.

Thus perhaps certain aspects of the experiment by Zajitschek *et al*. were not permissive of DR. These factors include diet (Simons & Dobson, 2023) and it is worth noting that the diets used by Zajitschek *et al*. are richer in yeast than those used here. Their standard diet contains the same amount of yeast as the richest diet used in our study. That a clear positive control, namely DR extending lifespan consistently, is missing further limits what we can interpret from this prior study.

Our work here, supported by prior published work by others (Kim et al., 2020; Shaposhnikov et al., 2022), does not support the interpretation that DR is a lab artefact and that DR would therefore not extend lifespan in the wild (Zajitschek et al., 2023). Even though conditions, environmental or genetic, that modulate the DR response are a valuable tool forward to understanding the DR response and ageing more generally (Simons & Dobson, 2023), it deters progress if such contrary findings are overinterpreted, especially when they are interpreted to have a broader ecological, evolutionary and physiological relevance.

## Methods

### Fly Husbandry and Diets

The ywR lab strain of *Drosophila melanogaster* was used for all experiments (Wessells et al., 2004). All flies were cultured on rich yeast media ([8% autolyzed yeast, 13% table sugar, 6% cornmeal, 1% agar, nipagin 0.225% (w/v), propanoic acid 0.4% (v/v)]. Cooked fly media was stored for up to 2 weeks at 4– 6°C, and warmed to 25°C before use. For all experimental diets (to which no propanoic acid was added), all components of the fly food media remained consistent but the amount of yeast was varied (2, 4, 6, 8 and 10% yeast), representing a spectrum of rich to restricted diets (McCracken, Buckle, et al., 2020; Simons & Dobson, 2023). For experimental diets, 8% yeast is the standard *ad libitum* diet used by the lab and 2% is the standard restricted diet used.

### Demography Protocol

Bottles of 10-12 females and 3-4 males per bottle were set up, allowing for 2 days egg laying before flies were removed to control growing density (Linford et al., 2013). The F1 generation from these bottles were transferred to new mating bottles as they eclosed and left to mate for 2 days. Newly eclosed offspring were transferred every day to generate age matched cohorts. After mating, offspring were anaesthetised using CO2 (Flystuff Flowbuddy; <5 L/min), females were sorted into groups of 70-100 and put into purpose-built demography cages (Good & Tatar, 2001; McCracken, Adams, et al., 2020), in which experimental diets were started. The cage design allowed for easy removal of dead flies and changing of fly food vials with minimal disruption to the population of flies. Every other day, food vials were replaced for each cage and a census of the flies was conducted. Any dead flies were counted and removed from the cage. Any escaped flies, or flies stuck to the fly food were right censored. Once sorted into cages, flies were housed in temperature-controlled incubators with humidity provided by large trays of water at either 21°C or 18°C (∼60% humidity, 12:12 light-dark cycle). For experiments conducted at 25°C, cages were kept in a climate-controlled room (60% humidity, 12:12 light-dark cycle) and the same census protocol was followed.

### Egg counting

Food vials were taken from demography cages 36 and 40 days after the experiments started, ages determined to be roughly a midpoint in the lifespan of the fly populations at the highest yeast diets. Eggs laid in the vials, constituting two days of egg laying, were counted manually under a light microscope.

### Data Analysis

Lifespan data were analysed using time-to-event mixed-effects Cox proportional hazard models, with demography cage as a random term (R package: coxme; function: coxme), to correct for uncertainty of pseudo replicated effects within cages (Therneau et al., 2003). The interaction between temperature and diet was fitted to test for differential effects of diet on mortality depending on temperature. Egg laying was analysed using linear models for each measurement time point separately. Egg counts were divided by the total females in the cage at the time of fecundity measurement to correct for differences in mortality.

## Acknowledgements

We thank Laura Hartshorne and Gracie Adams for helping run the experiments.

